# Transient glycolytic complexation of arsenate enhances resistance in the enteropathogen *Vibrio cholerae*

**DOI:** 10.1101/2022.08.04.502822

**Authors:** Emilio Bueno, Víctor Pinedo, Dhananjay D. Shinde, André Mateus, Athanasios Typas, Mikhail M Savitski, Vinai C. Thomas, Felipe Cava

## Abstract

The ubiquitous presence of toxic arsenate (As^V^) in the environment has virtually raised mechanisms of resistance in all living organisms. Generally, bacterial detoxification of As^V^ relies on its reduction to arsenite (As^III^) by ArsC, followed by the export of As^III^ by ArsB. However, how pathogenic species resist this metalloid remains largely unknown. Here, we found that *V. cholerae,* the etiologic agent of the diarrheal disease cholera, outcompetes other enteropathogens when grown on millimolar concentrations of As^V^. To do so, *V. cholerae* uses, instead of ArsCB, the As^V^-inducible *vc1068-1071* operon (renamed *var* for *v*ibrio *a*rsenate *r*esistance), which encodes the arsenate repressor ArsR, an alternative glyceraldehyde-3-phosphate dehydrogenase, a putative phosphatase, and the As^V^ transporter ArsJ. In addition to Var, *V. cholerae* induces oxidative stress- related systems to counter ROS production caused by intracellular As^V^. Characterization of the *var* mutants suggested these proteins function independently from one another and play critical roles in preventing deleterious effects on the cell membrane potential and growth derived from the accumulation As^V^. Mechanistically, we demonstrate that *V. cholerae* complexes As^V^ with the glycolytic intermediate 3-phosphoglycerate into 1-arseno-3-phosphoglycerate (1As3PG). We further show that 1As3PG is not transported outside the cell; instead, it is subsequently dissociated to enable extrusion of free As^V^ through ArsJ. Collectively, we propose the formation of 1As3PG as a transient metabolic storage of As^V^ to curb the noxious effect of free As^V^. This study advances our understanding of As^V^ resistance in bacteria and underscores new points of vulnerability that might be an attractive target for antimicrobial interventions.

## Introduction

Arsenic is a toxic metalloid commonly found in aquatic and terrestrial environments (1–3) as arsenate (As^V^) or its reduced form, arsenite (As^III^). The potent toxicity of this element makes it one of the best-studied natural poisons that impact public health (4). The mechanism of As^V^ toxicity is due to its structural resemblance to phosphate oxyanions and its uptake into the cell through phosphate transport channels (5, 6). Once in the cytoplasm, As^V^ can replace phosphate in energy-generating reactions resulting in the formation of ADP-As^V^ instead of ATP by the ATPase, and through substrate-level phosphorylation during glycolysis (7, 8). Conversely, cellular toxicity by As^III^ is exerted by a different mechanism. As^III^ can interact directly with thiol groups of proteins and molecules, thereby interfering with diverse cellular processes (9, 10). Despite its risk to public health, arsenic has been used as an antimicrobial agent to treat infectious diseases (1, 4, 11, 12), increasing the selective pressure for microbes to acquire arsenic resistance.

Bacterial strategies to resist As^V^ include chelation by metal-binding metallothioneins (13), methylation to less toxic and more volatile arsenic derivatives (14), respiration (15), and extrusion of As^V^ or As^III^ by specific efflux pumps (e.g., ArsJ and ArsB, respectively) (9, 16–18). In bacteria, the most common mechanism of As^V^ resistance genes occurs via As^V^ reduction to As^III^ followed by the extrusion of the latter through an As^III^ transporter (19). As^V^ resistance genes are commonly organized in operons, e.g., the *arsRDABC* operon from *Escherichia coli* R773 (19). In this example, *arsC* encodes a dedicated arsenate reductase and *arsB* an efflux pump of As^III^ (20). The *arsR* gene encodes the repressor of the *ars* operon. When As^III^ is present, its interaction with ArsR releases it from the *ars* promoter, thus increasing transcription of the operon. *arsA* encodes an ATPase that, together with ArsB forms the As^III^ extrusion system. Alternatively, As^III^ extrusion via ArsB can be driven by proton motive force in species lacking ArsA. Finally, *arsD* encodes a chaperone that enhances arsenic extrusion by transferring As^III^ to ArsAB (9).

Despite being extensively studied in environmental soil and marine bacteria species, yet the mechanisms underlying As^V^ resistance in human pathogenic bacteria remain largely unknown. Here, we investigated As^V^ resistance in representative enteric pathogens and found that *V. cholerae* exhibits remarkably high resistance to this metalloid. Using genome mining and functional characterization, we demonstrated that *V. cholerae* does not detoxify As^V^ by reducing it to As^III^. Instead, transposon-based functional screens identified the operon *vc1068-1071* as the primary genetic determinant of As^V^ resistance in *V. cholerae*. Mechanistic characterization of this system revealed that resistance to As^V^ in *V. cholerae* is mediated by a transient metabolic complexation of As^V^ with the glycolytic intermediate 3-phosphoglycerate (3PG) to generate 1-arseno-3- phosphoglycerate (1As3PG). Our results support that this As^V^-containing metabolite is transient in the cell: its formation cushions the cellular damage caused by free As^V^, while its dissociation enables *V. cholerae* to release free As^V^ through the ArsJ efflux permease.

## Results

### High resistance to arsenate in *V. cholerae* is independent of the ArsC reductase

While studying the tolerance of enteropathogenic bacteria to metalloids we observed that *Vibrio cholerae*, the causative agent of cholera, was able to grow in media supplemented with supraphysiological (30 mM) concentrations of arsenate (As^V^) (Fig. S1). This level of resistance was unmatched by other human enteric pathogens, as lower concentrations of As^V^ (10 mM) readily caused total or partial growth inhibition of *Salmonella enterica*, *Citrobacter rodentium, Yersinia pseudotuberculosis, Enterohemorragic E. coli* (EHEC) *and Shigella flexneri* (Fig. 1A). Consistently, *V. cholerae* outcompeted these species by 5-500-fold in co- culture experiments grown in media supplemented with As^V^ (Fig. 1B).

**Fig. 1.**
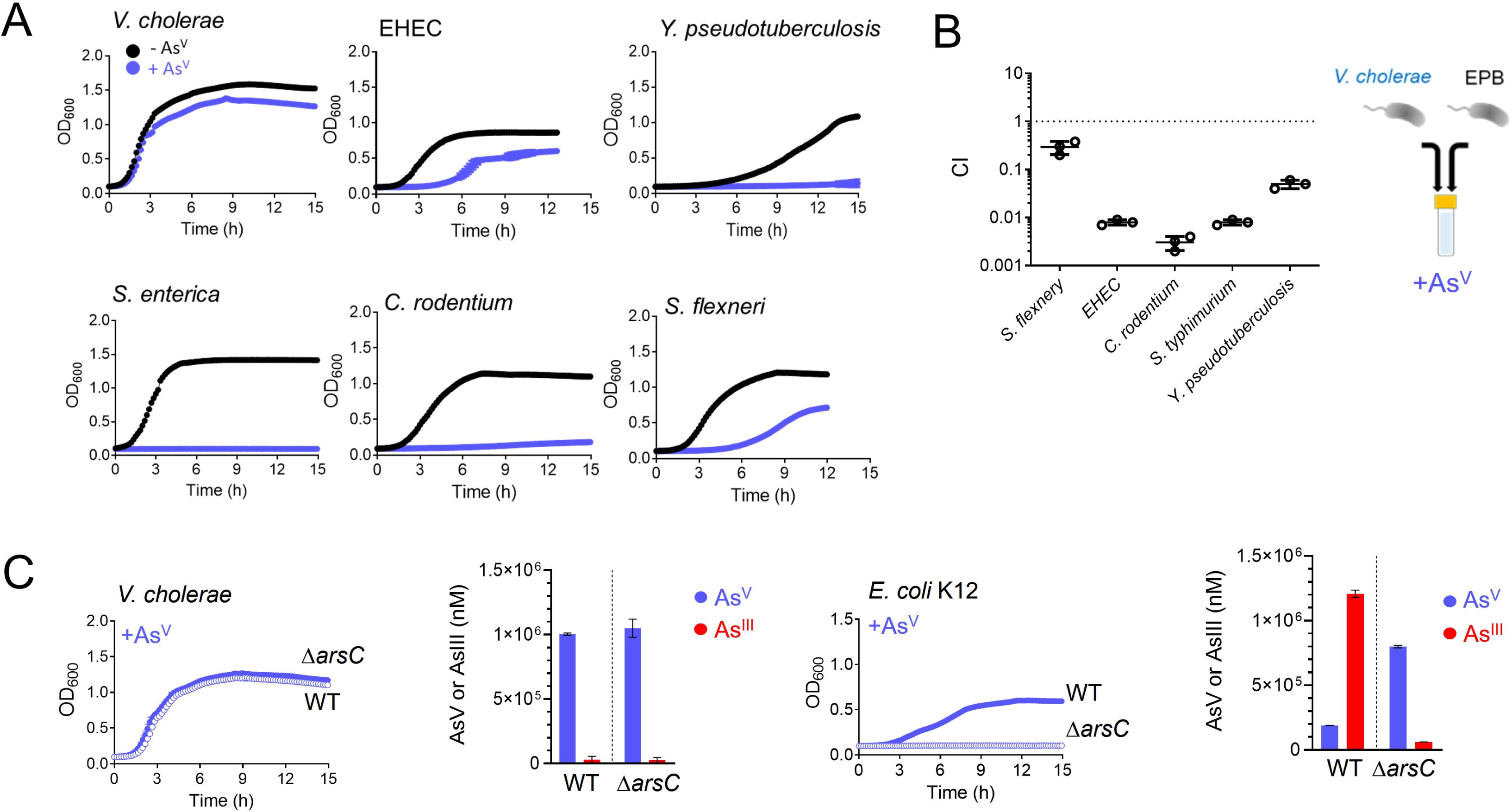
*V. cholerae* is the enteric pathogen with the greatest resistance to As^V^. **A** Growth curves (OD_600_) of the enteric pathogens *V. cholerae*, *Enterohaemorragic Escherichia coli* (EHEC), *Salmonella enterica*, *Citrobacter rodentium*, *Shigella flexneri*, and *Yersinia pseudotuberculosis*. Cultures were grown in LB medium in the absence (black) and in the presence (blue) of 10 mM arsenate (As^V^). **B** *in vitro* competition among *V. cholerae* and other enteropathogenic bacteria (EPB) studied in panel A. Cells were incubated in LB medium supplemented with 10 mM As^V^ for 8 hours. CI values from competition in the presence of As^V^ were normalized with respect to CI values in the absence As^V^. **C** Growth curves (OD_600_) of WT (blue filled circles) and *arsC* (blue empty circles) mutant strains from *V. cholerae* and *E. coli* K12. Cultures were grown in LB medium supplemented with 10 mM As^V^. As^V^ and As^III^ concentrations were determined by ICP-MS from supernatants of *V. cholerae* and *E. coli* WT and *arsC* mutant strains after 8 hours of incubation in 1.5 mM of As^V^. Data are the mean of three biological replicatesD±Ds.e.m.

To investigate the molecular mechanisms of As^V^ resistance in *V. cholerae*, we first searched for canonical determinants for As^V^ detoxification in its genome. *V. cholerae vc2165* encodes a homolog of *E. coli* K12 arsenate reductase ArsC, including the conserved catalytic cysteine and arginine residues (36, 37) (Fig. S2A). However, while *E. coli*’s ArsC enzyme is critical for resistance to As^V^, inactivation of *vc2165* did not compromise *V. cholerae* growth in the presence of this metalloid (Fig 1C). Thus, we reasoned that *V. cholerae*’s ArsC might not be active or expressed. To measure ArsC activity we quantified extracellular As^V^ and As^III^ by inductively coupled plasma mass spectrometry (ICP-MS) and observed that while *E. coli* reduced ∼80% of As^V^ to As^III^ in an ArsC-dependent manner, *V. cholerae* produced no As^III^ (Fig. 1C). Interestingly, replacement of *V. cholerae arsC* allele by that of *E. coli* caused growth inhibition, likely due to toxic As^III^ formation, as growth was recovered by expressing *E. coli’s* As^III^-transporter ArsB (Fig. S2B). Altogether, these results indicate that *V. cholerae* ArsC is inactive and that As^V^ resistance in this bacterium is independent of the ArsC-AsrB system that relies on the production and elimination of As^III^.

### Resistance to As^V^ in *V. cholerae* depends on the *vc1068*-*vc1071* operon

To identify the genetic determinants conferring resistance to As^V^ in *V. cholerae*, we performed two complementary genome-wide screenings based on: ***i*)** Transposon Insertion Sequencing (TIS), and ***ii*)** the use of a previously described *V. cholerae* transposon mutant library [32] (Fig. 2). Both screens permit assessment of the mutants’ relative fitness. However, the first approach is more suitable for discovering mutants with subtle fitness differences, while the latter can identify candidates otherwise underappreciated by TIS due to cross- complementation between mutants within a population. Only four non-essential Tn mutants (*vc1068::tn, vc1069::tn, vc1070::tn,* and *vc1071::tn*) were mutually under-represented in both screenings (Fig. 2). We renamed *vc1068-71* as ***var*** for ***v****ibrio* **a**rsenate **r**esistance cluster referred hereafter for reasons described below. *vc1068* encodes an homolog of the ArsR repressor, *vc1069* (renamed as *varG*) encodes a putative glyceraldehyde-3-phosphate dehydrogenase (GAPDH), *vc1070* (renamed as *varH*) encodes an uncharacterized putative phosphatase, and *vc1071*, an ArsJ-like As^V^ transporter (17). As expected, the *arsC* mutant was not identified in the screens, confirming that *V. cholerae* does not detoxify As^V^ by reducing it to As^III^. Although homologues to VarG and ArsJ were previously reported in *Pseudomonas aeruginosa*, their function in As^V^ resistance has not been studied *in vivo* (17).

**Fig. 2.**
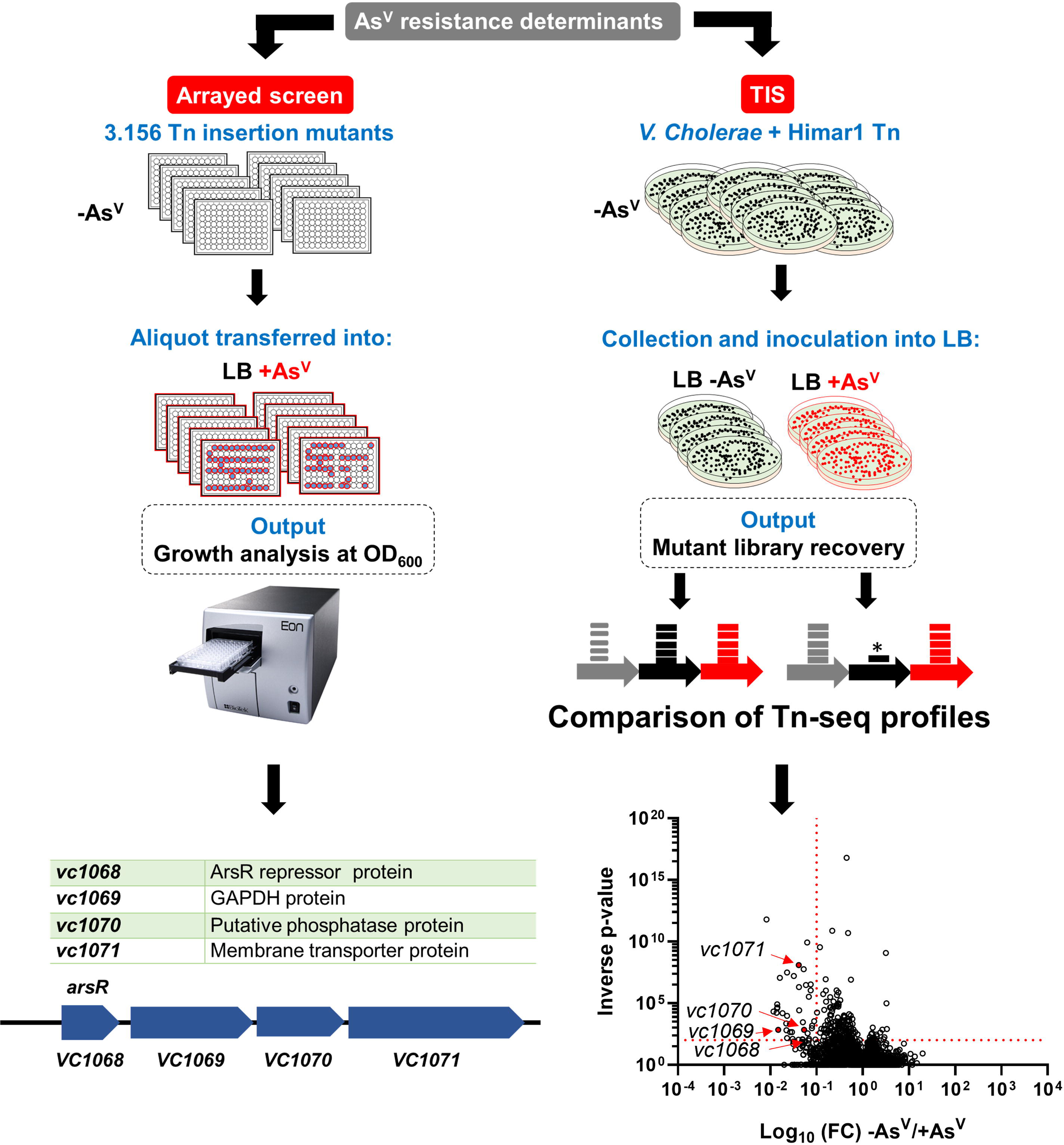
Identification of essential genetic determinants for As^V^ resistance in *V. cholerae*. Left: Experimental workflow for an arrayed transposon mutant screen to identify the genes required for As^V^ resistance in *V. cholerae* and the list of mutant strains identified. Right: Experimental workflow for TIS screen in the presence and absence of 1 mM As^V^. Volcano plot depicting the ratio of read counts mapped to individual genes in transposon libraries of *V. cholerae* plated onto media supplemented with As^V^). Red dotted lines indicate arbitrary thresholds of fold change (FC) <0.1 and an inverse p-value >20. Genes in red color show loci also identified in the arrayed screen (see Supplementary Table 1 for an extended version).

To verify the implication of the *var* cluster on As^V^ resistance, we constructed individual in-frame marker-less deletion mutants. All mutants were sensitive to As^V^except for Δ*arsR* (Fig. 3A), suggesting that the conditional lethality of the *arsR::*tn was likely due to the polar effect on the expression of the downstream genes *varG- varH-arsJ*. Indeed, transcriptional analysis using fusions of the *lacZ* reporter to potential promoter regions located upstream of each *var* gene demonstrated that expression of this cluster depends on As^V,^ and it is driven from a single promoter upstream of *arsR* (Fig. S3). Inactivation of *arsR* turned transcription from this promoter constitutive (both in presence and absence of As^V^), confirming that VC1068 is the *V. cholerae*’s ArsR repressor of the *var* operon (Fig. S3). Further characterization revealed that Δ*arsJ* presents a more dramatic growth defect compared to the rest of the *var* mutants at low As^V^ concentrations (1 mM) (Fig. 3). However, at 10 mM of As^V^, the growth of the *varG* and *varH* mutant strains was also compromised despite the presence of ArsJ. These results suggest that the ArsJ transporter is the main determinant for As^V^ resistance in *V. cholerae* but VarG and VarH also become relevant at higher concentrations of As^V^. Furthermore, the inactivation of *varG, varH* and *arsJ* (double and triple mutants) aggravated growth defects exhibited by the single mutants, suggesting independent functions for these proteins in As^V^ resistance in *V. cholerae* (Fig. 3A).

**Fig. 3.**
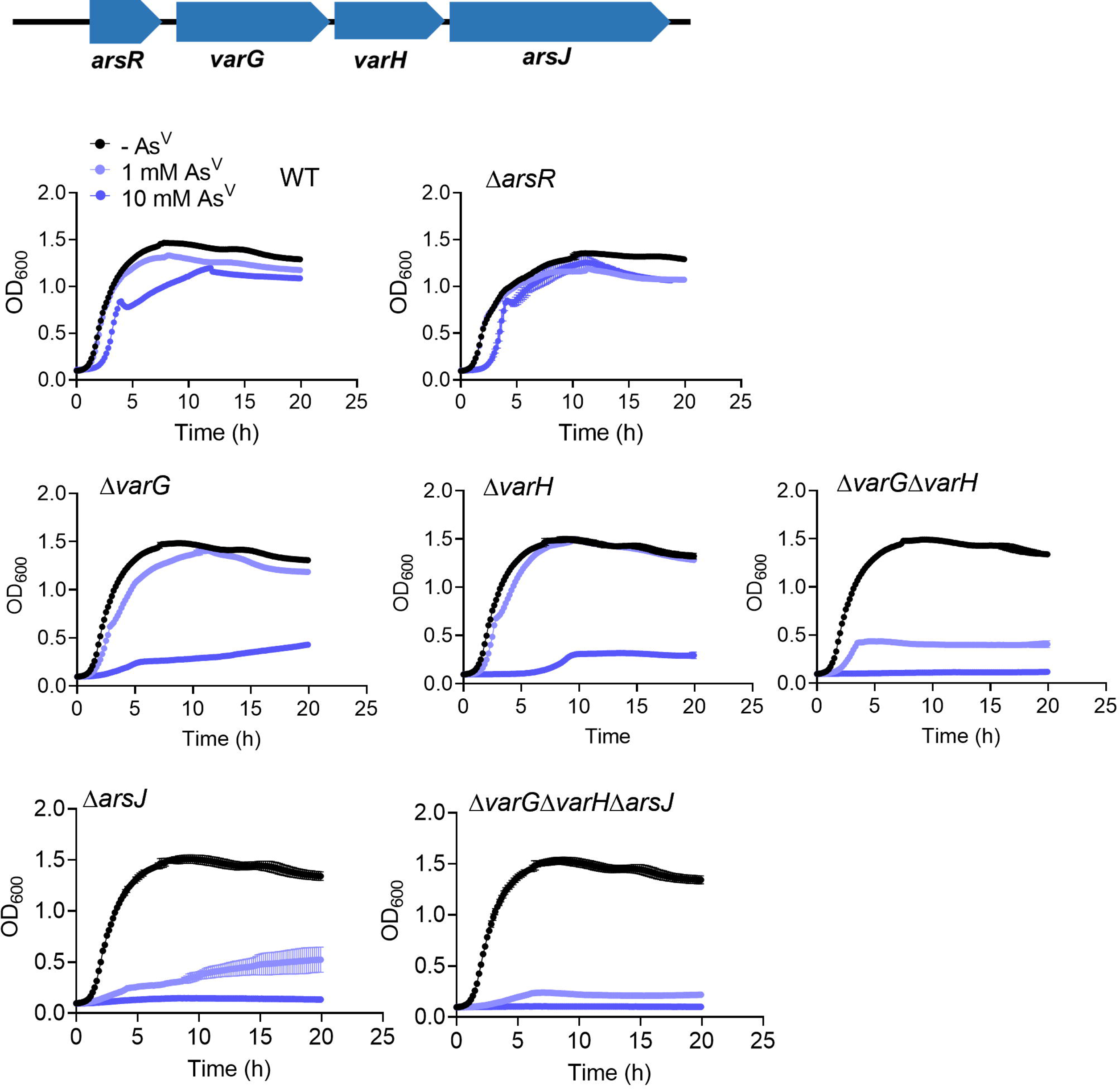
Phenotypic characterization of the *V. cholerae var* operon. Growth curves (OD_600_) of *V. cholerae* WT, single and combinatorial *var* mutant strains. Cultures were grown in LB medium in the absence or presence of 1 and 10 mM As^V^. Data are the mean of three biological replicatesD±Ds.e.m.

### VarG is an As^V^-inducible GAPDH that preferentially binds As^V^

*V. cholerae* VarG presents a high degree of protein sequence (E-value: 5^e-86^) (Fig. S4A) and structure (AlphaFold2 TM-score=0.96) (Fig. S4B) similarity to the glycolytic GAPDH VC2000 (Gap), including the conserved catalytic Cys residue (38). To characterize VarG’s activity compared to the glycolytic paralogue Gap particularly, we purified both proteins from *V. cholerae* and performed *in vitro* GAPDH activity assays (39) in the presence or absence of As^V^. While both proteins exhibited GAPDH activity, VarG showed preference for As^V^ over P_i_ (the canonical substrate) (Fig. 4A). Conversely, the glycolytic GAPDH Gap performed better with P_i_. Furthermore, while the expression of *Gap* remained low and constitutive, transcription from the *var* promoter was strongly induced by As^V^ (Fig. 4B). Therefore, these results suggest that while *V. cholerae* Gap is active on As^V^, this enzyme specific activity and protein levels are likely insufficient to replace VarG essentiality in *V. cholerae* resistance to As^V^.

**Fig. 4.**
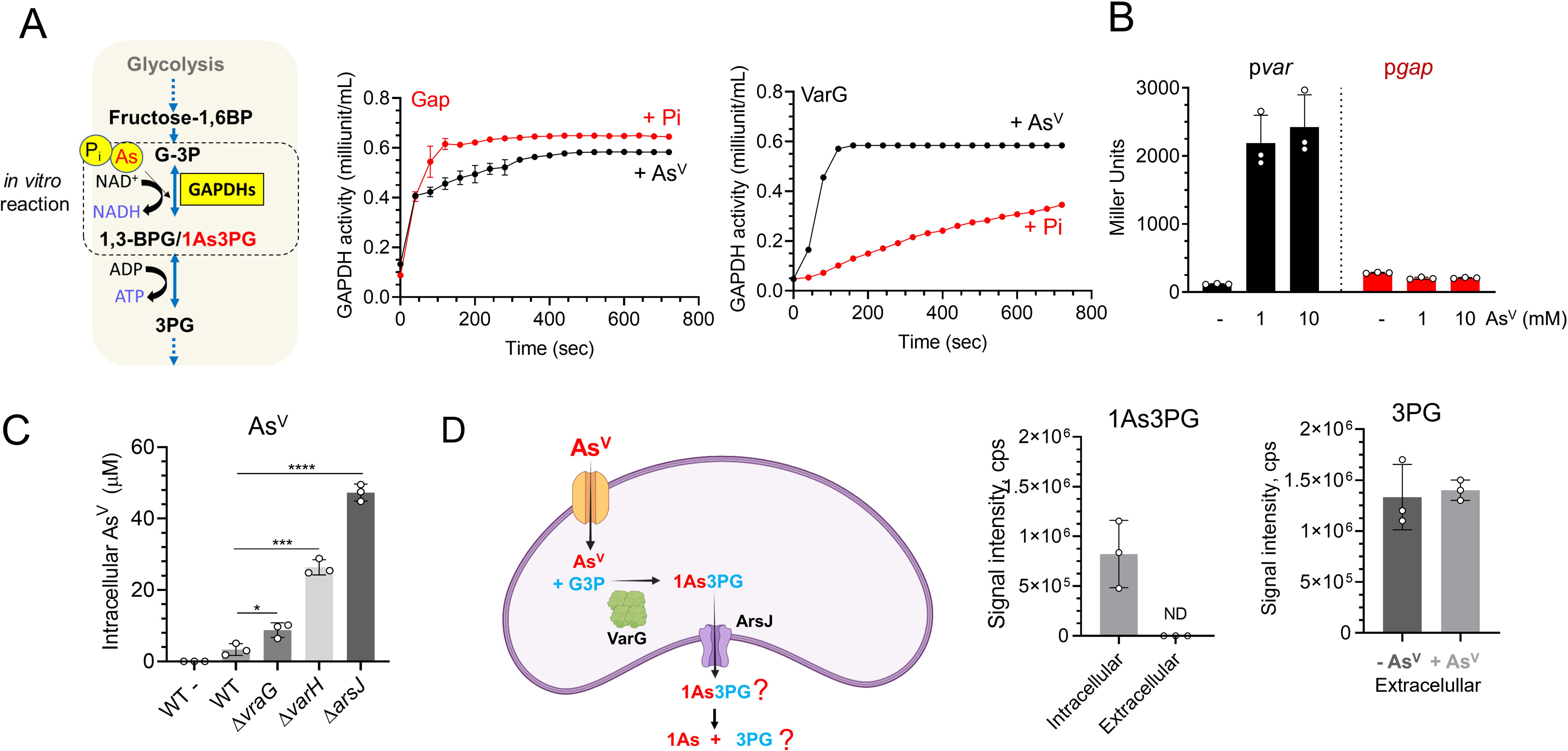
Characterization of *V. cholerae* VarG. A *in vitro* GAPDH activity (OD_450_) of purified *V. cholerae* VarG and the glycolytic Gap enzymes using P_i_ or As^V^ as substrates. **B** β-galactosidase activity of strains in which the *lacZ* reporter gene is cloned under the control of the *var* operon and the *gap* genes promoters. Cultures were grown in LB medium in the absence or presence of 1 and 10 mM As^V^ for 5 hours. **C** Intracellular As^V^ concentrations of WT, *varG, varH* and *arsJ* mutant strains grown with 1 mM As^V^by ICP-MS. **D** Left: As^V^ detoxification model in *P. aeruginosa* DK2 [1]. Right: Quantification of 1As3PG and 3PG from intracellular and extracellular samples of *V. cholerae* cells grown with and without 1 mM As^V^. Data are the mean of three biological replicatesD±Ds.e.m.

### Arsenate detoxification in *V. cholerae* is facilitated by extrusion of free arsenate instead of complexed as 1-arseno-3-phosphoglycerate

A VarG homolog from *P. aeruginosa* was proposed to function as a GAPDH that complexes As^V^ with glyceraldehyde-3-phosphate (G3P) into 1-arseno-3- phosphoglycerate (1As3PG), which is further eliminated through ArsJ (17). However, these experiments were performed *in vitro* using a commercial glycolytic GAPDH from rabbit instead of the As^V^-specific GAPDH from *P. aeruginosa* DK2. If this model applies to *V. cholerae,* the absence of either VarG or ArsJ will similarly affect AsV export and resistance. However, our data shows that Δ*varG* did not phenocopy the growth defect of Δ*arsJ* (Fig. 3). Furthermore, analysis of intracellular levels of As^V^ by ICP-MS revealed that, in relation to the WT, while the Δ*varG* strain only showed moderate (ca. 3x) accumulation of As^V^, inactivation of ArsJ accumulated this metalloid ca. 20x (Fig. 4C). Although these results suggest that VarG is not essential to export As^V^, we cannot discard the possibility of As^V^ being complexed and extruded as 1As3PG in *V. cholerae*.

Using high-resolution mass spectrometry (HRMS), we detected a mass (*m/z*) of 304.8696 Da, specific to the cultures supplemented with As^V^, which was compatible with 1As3PG (*m/z* 304.8674, *m/z*<0.003 Da difference between experimental and theoretical mass) (Fig. 4D and S5). 1As3PG was also detected in the Δ*varG*, consistent with the ability of the constitutively expressed glycolytic Gap GAPDH to use As^V^ (Fig. S5). 1As3PG was detected in intracellular cytoplasmic fractions but not in the extracellular milieu, suggesting that this molecule is not expelled in *V. cholerae* (Fig. 4D and S5). As it has been reported that 1As3PG is rapidly dissociated to As^V^ and 3PG (40), we reasoned that if exported, its decomposition should increase extracellular 3PG levels in relation to cultures without As^V^. However, we found no significant differences in the levels of extracellular 3PG between conditions supporting the idea that As^V^ detoxification likely occurs through the release of free As^V^ rather than of 1As3PG through ArsJ (Fig. 4C and S5).

### VarH is involved in 1As3PG dissociation and arsenate export

To find clues that help us understand the role that 1As3PG plays in *V. cholerae*’s resistance to As^V^ we investigated VarH. Structural *in silico* analysis using the phyre2 tool (www.sbg.bio.ic.ac.uk/phyre2) and AlphaFold2 revealed that VarH is a cysteine-based protein tyrosine phosphatase (PTP) belonging to the dual- specificity phosphatase superfamily that presents a very high topology similarity to the human Kap PTP phosphatase (Fig. S6AB). To determine if VarH has phosphatase activity, we purified this protein and assessed its capacity to dephosphorylate the chromogenic substrate pNPP. Our results show that, compared to the wheat phosphatase or *V. cholerae’s* PTP VC1041 positive controls, VarH presents lower phosphatase activity, which turned off upon replacement of the putative catalytic cysteine and arginine by glycine residues (Fig. 5A, left panel). Furthermore, consistent with the importance of these residues in VarH activity, replacement of the WT *varH* allele by the catalytically inactive variant *varH* C113G/R119G impaired *V. cholerae* growth in the presence of As^V^ (Fig. 5A, right panel). These results suggest that the phosphatase activity of VarH is required to provide As^V^ resistance in *V. cholerae*.

**Fig. 5.**
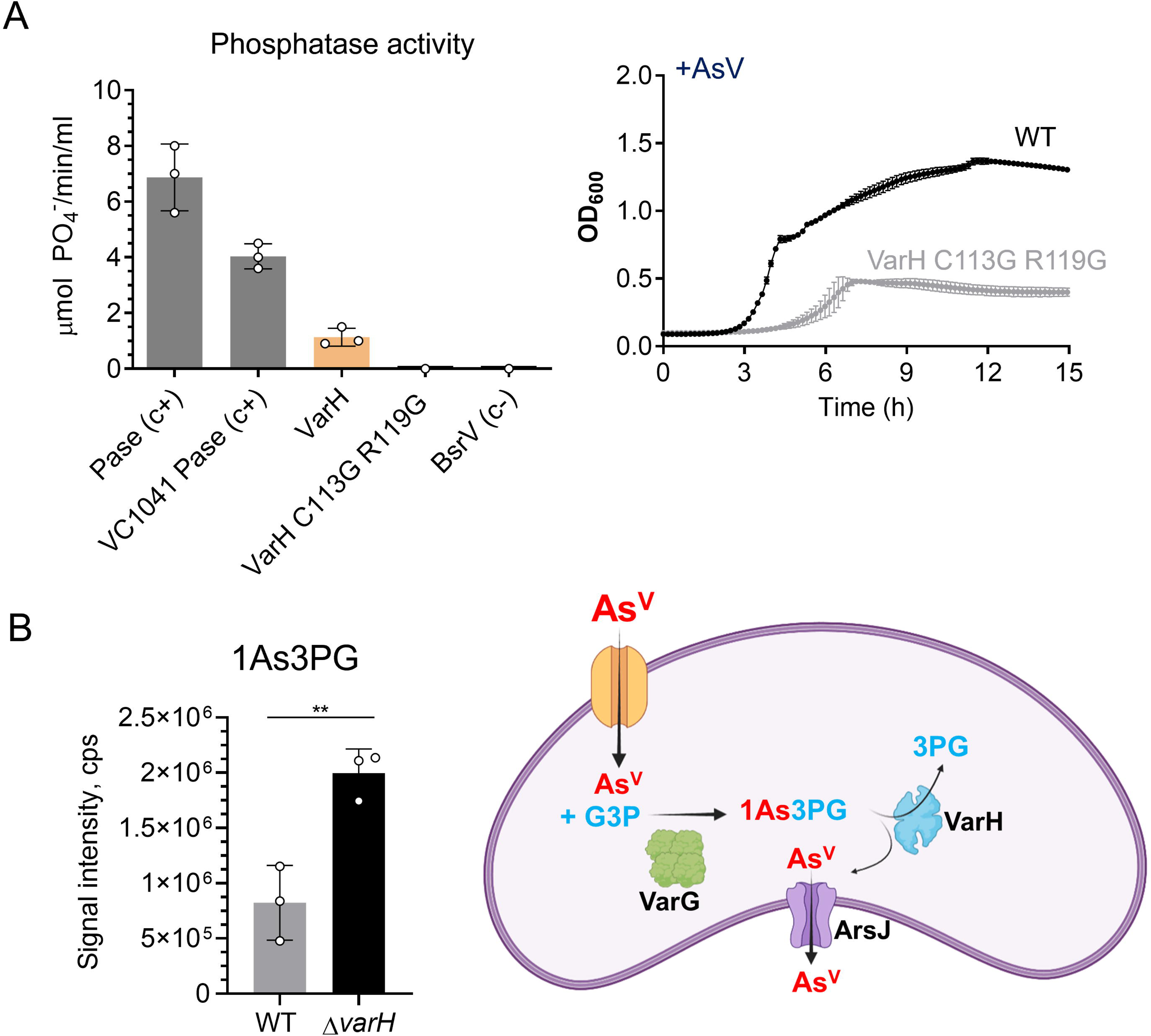
Characterization of *V. cholerae* VarH. **A** Left: *in vitro* phosphatase activity of purified *V. cholerae* VarH, and catalytic mutant derivative where cysteine 113 and arginine 119 were replaced by glycine. *V. cholerae* protein- tyrosine-phosphatase (VC1041) and broad-spectrum racemase (BsrV) were used as positive and negative controls, respectively. Right: Growth curves (OD_600_) of *V. cholerae* WT, and a VarH catalytic mutant derivative . Cultures were grown in LB medium in the presence of As^V^. **B** Left: Intracellular 1As3PG concentrations from *V. cholerae* WT and Δ*varH* cells grown with and without 1 mM As^V^ by HRMS. Right: Schematic of putative mechanism for As^V^ detoxification in *V. cholerae*. Data are the mean of three biological replicatesD±Ds.e.m.

Since VarG and VarH are coregulated by As^V^ and both support resistance to this metalloid, we hypothesized that VarH could regulate VarG activity by modulating its phosphorylation state. To assess such a possibility, we purified VarG from *V. cholerae* WT and Δ*varH* (Fig. S6C) and analyzed VarG phosphorylation by liquid chromatography–tandem mass spectrometry (LC– MS/MS). However, even though VarG was phosphorylated on Tyr108 and Ser204, those phosphorylation levels were independent of VarH (Fig. S6D). Then, we reasoned that as P_i_ and As^V^ are analogs, VarH could potentially act on 1As3PG as a phosphatase to produce 1AsG + Pi or as an “arsenatase” to render 3PG + As^V^. Remarkably, we found that 1As3PG concentration were significantly elevated in Δ*varH* compared to those in the WT strain (Fig. S6E), suggesting that VarH might use 1As3PG as a substrate. Although we could not confirm VarH’s activity *in vitro,* potentially due to the instability of 1As3PG (41), the fact VarH is an active phosphatase but 1AsG is not detected in *V. cholerae* suggests that VarH could dissociate the 1As3PG complex into 3PG and free As^V^ which would be released through ArsJ.

### Intracellular arsenate impacts the *V. cholerae* proteome and leads to ROS accumulation and membrane potential defects

To broadly assess global responses triggered by As^V^ in *V. cholerae*, we analyzed the pathogen proteome upon exposure to As^V^. Interestingly, a relatively low number of proteins (D20 proteins) were upregulated upon exposure to As^V^, indicating that adaptations to this metalloid might not require extensive proteome rewiring in *V. cholerae*. As expected, these proteins included the As^V^ resistance Var proteins but also amino acids permeases, a Zn/Cd transporter, members of the PTS-fructose system, the starvation stress response protein RaiA, and the cysteine synthesis and the peroxiredoxin PrxA (Fig. 6AB). Cysteine is one of the main targets of reactive oxygen species (ROS) and a functional component of glutathione, which plays a crucial role in the adjustment of cellular redox potential (42). In this line, PrxA has been involved in the adaptation to hydrogen peroxide in *V. cholerae* (43). Therefore, we reasoned that the *var* mutants, which display increased As^V^ intracellular levels, might suffer from higher oxidative stress than *V. cholerae* WT strain. To test this hypothesis, we measured ROS levels in the *V. cholerae* WT and *var* mutant strains and observed a direct correlation between As^V^ accumulation and ROS level (Δ*arsJ*>Δ*varH*>Δ*varG*>WT) (Fig. 6C). These findings collectively indicate that AsV induces oxidative stress in *V. cholerae* which is exacerbated in the *var* mutants due to a supraphysiological accumulation of this toxic metalloid.

**Fig. 6.**
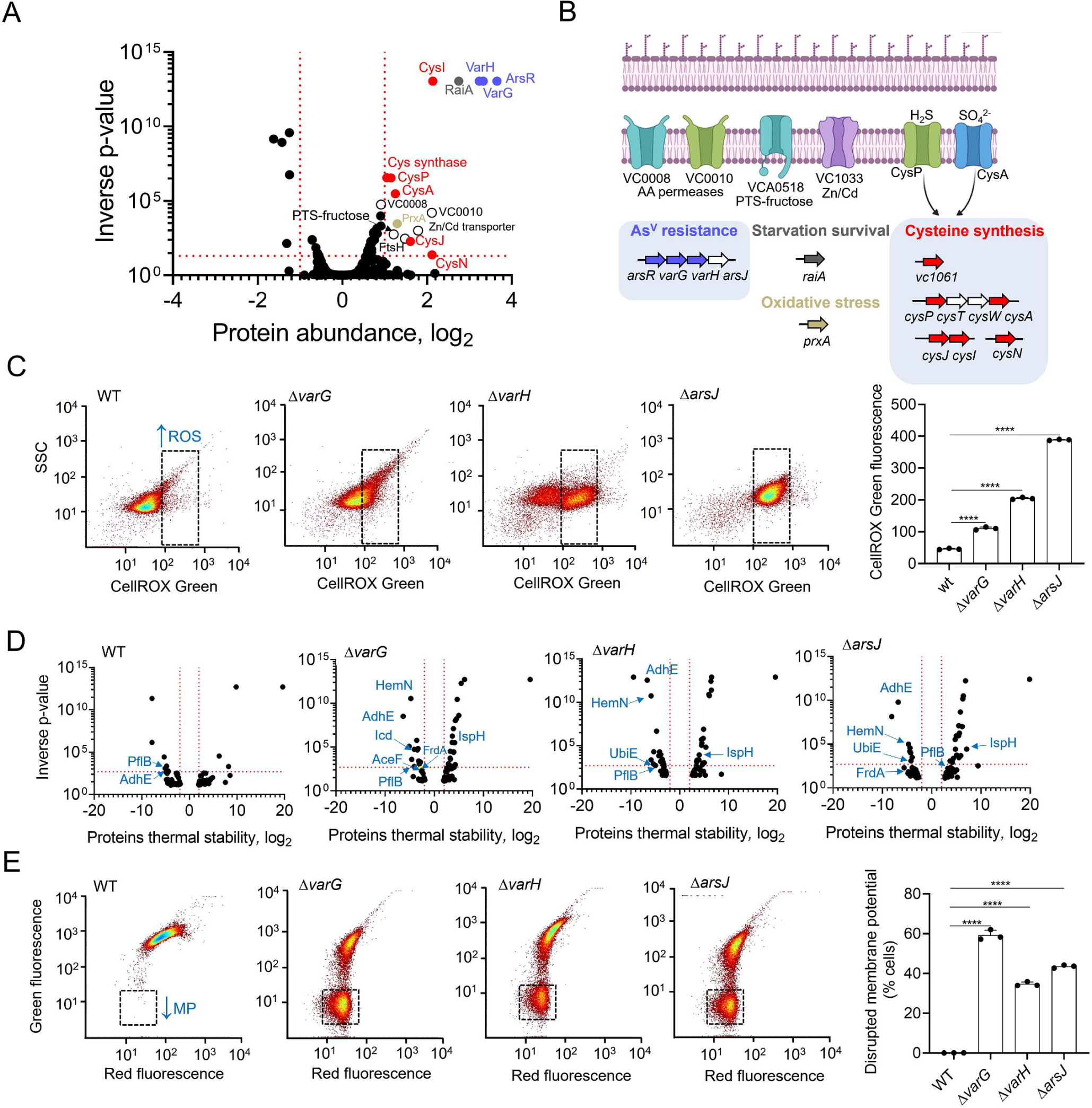
Oxidative stress response and membrane potential alterations of *V. cholerae* WT and the *var* mutants. **A** Volcano plot of LIMMA analysis output depicting read fold change (log_2_) of protein abundance and inverse P-value for each protein queried in the proteomic screen in response to 1 mM As^V^ in *V. cholerae* WT. Red dotted lines indicate arbitrary thresholds of fold change (log_2_) of >1 or <1 and an inverse P-value of >20. Var proteins are labelled in blue and oxidative stress related proteins (Cystein biosynthesis and PrxA are labelled as in B. **B** Schematic of the systems induced in response to As^V^ in *V. cholerae*. Non- coloured arrows depict genes within an operon not detected. **C** Representative dot plots from *V. cholerae* WT and *var* mutant cells stained with CellRox Green following growth in the presence of 1 mM As^V^ for 5 hours. The black dotted gate indicates an area with higher CellRox fluorescence (higher ROS levels). **D** Volcano plot of LIMMA analysis output depicting read fold change (log_2_) of protein thermal stability and inverse P value for each protein queried in the proteomic screen in response to 1 mM As^V^ in *V. cholerae* WT and *var* mutant strains. As^V^ interacting proteins related to energy-generating pathways are labeled in blue color. PflB, VC1866 formate acetyltransferase. AdhE, VC2033 alcohol dehydrogenase/acetaldehyde dehydrogenase. HemN, VC0116 oxygen- independent coproporphyrinogen III oxidase. Icd, VC1141 isocitrate dehydrogenase. FrdA, VC2656 fumarate reductase. IspH, 4-hydroxy-3-methylbut- 2-enyl diphosphate reductase. AceF, VC2413 pyruvate dehydrogenase, E2 component. UbiE, VC0083 ubiquinone/menaquinone biosynthesis methyltransferase. **E** Representative dot plots from *V. cholerae* WT and *var* mutant cells stained with DiOC2 following growth in the presence of 1 mM As^V^ for 5 hours. The black dotted gate indicates an area with lowered green fluorescence (lower membrane potential). Data are the mean of three biological replicatesD±Ds.e.m.

In addition to ROS affecting Cys-containing proteins, we cannot discard a direct effect of As^V^ on the proteome of *V. cholerae*. To this end, we performed thermal proteome profiling (TPP), a method that globally analyzes multiple types of protein interactions (25). TPP data showed an increased thermal stability of multiple cytosolic proteins that could potentially interact with As^V^ (D3-fold) in the *var* mutants with respect to the WT (Fig. 6D and table S). As^V^-interacting protein candidates included proteins involved in the synthesis of heme groups of respiratory cytochromes, respiratory ubiquinone, fermentation, and carbon catabolism, suggesting that accumulation of As^V^ in Var-defective strains might interfere with cell bioenergetics. To investigate this possibility, we monitored the bacterial membrane potential in the presence of As^V^ using flow cytometry (Fig. 6E). Interestingly, inactivation of the *var* system resulted in a significant depolarization of the mutant’s membrane. Together, these results demonstrate that AsV can, directly and indirectly, affect the *V. cholerae*’s proteome, leading to oxidative stress and defects in cellular bioenergetics, both possibly related to the growth impairment of the *var* mutants.

### *V. cholerae var operon* provides As^V^ resistance to enteric pathogens

*In silico* analysis showed that, except for *C. rodentium*, ArsR is the only As^V^- resistance determinant conserved between *V. cholerae* and the As^V^-sensitive enteric pathogens (Fig. 7A). In general, these enteropathogenic species detoxify As^V^ via ArsC-dependent reduction to As^III^, and subsequent extrusion through ArsB. As *V. cholerae* lacks the As^III^- production/extrusion tandem ArsCB, we wondered whether the *var* genes could also augment resistance to As^V^ in other species or would rather require additional species-specific elements. To address this question, we expressed the *var* operon excluding the ArsR repressor in enteropathogens sensitive to As^V^ and studied their growth capacity in the presence of As^V^. Strikingly, induced coexpression of *varG-varH-arsJ* in several enteropathogens boosted their growth on As^V^ (Fig. 7B), demonstrating that the Var system is sufficient and compatible with the ArsCB system in providing resistance against As^V^ in other bacteria.

**Fig. 7.**
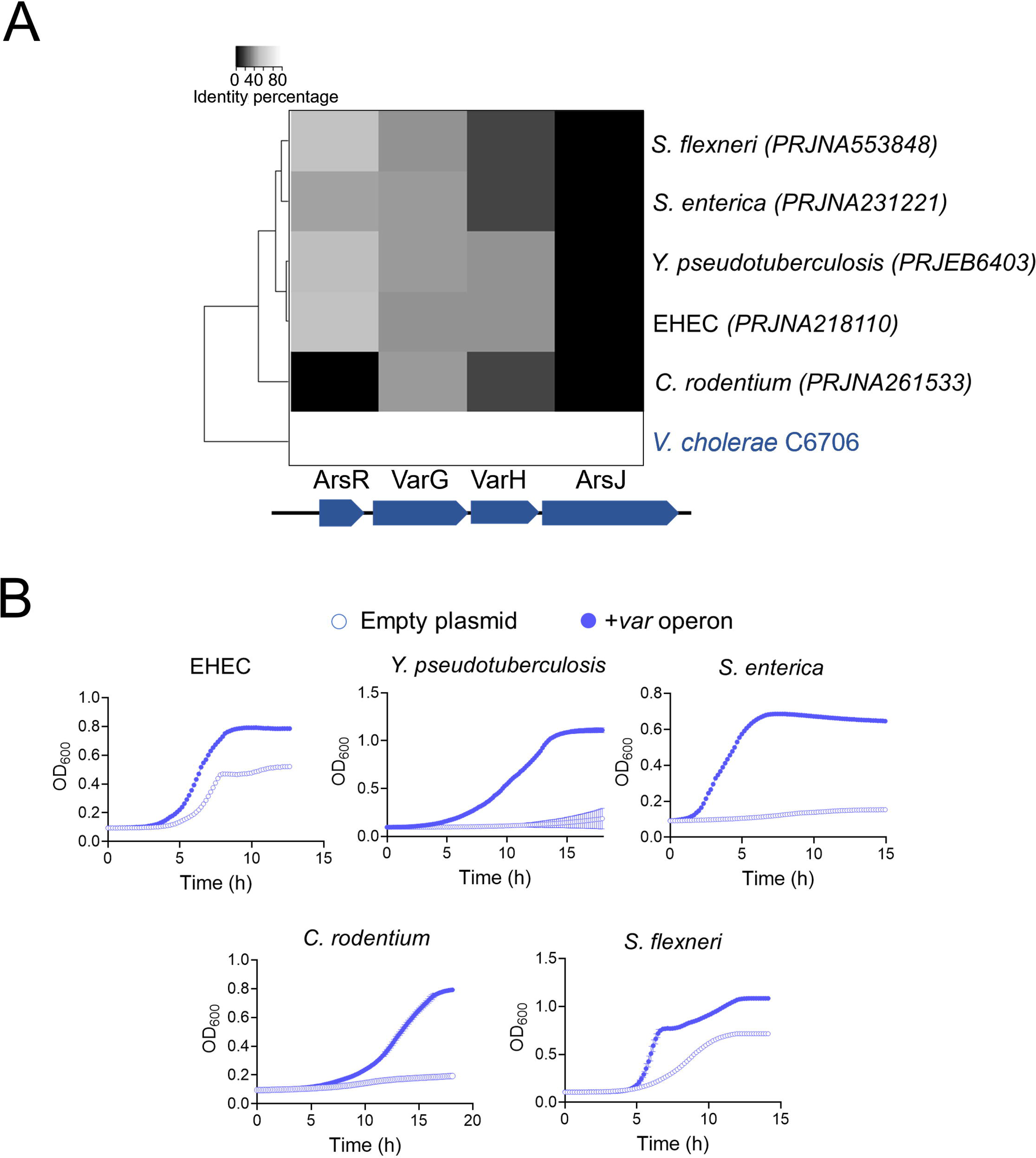
*V. cholerae* Var enhances As^V^ resistance in enteropathogenic bacteria. **A** Hierarchical heat map clustering of enteropathogenic bacteria with respect to *V. cholerae* based on the percentage identity of *var* genes . **B** Growth curves (OD_600_) of the As^V^-sensitive enteropathogenic bacteria clustered in panel A, carrying an inducible empty plasmid (empty circle) or the same plasmid expressingthe *V. cholerae var* operon (blue filled circle). Cultures were grown in LB medium in the presence of 10 mM As^V^. Data from panel B are the mean of three biological replicatesD±Ds.e.m.

## DISCUSSION

Arsenic is ubiquitous in the environment and consequently, most living organisms, including bacteria, encode arsenic-resistance proteins (44–49). However, functional characterization of these systems and how they support bacterial fitness in environments with As^V^ is underappreciated. In this study, we describe that *V. cholerae* exhibits higher resistance to As^V^ than other enteric pathogens.

The strategy used by *V. cholerae* to resist As^V^ is different from that used by other gammaproteobacterial enteropathogens. While most of these species encode an ArsC arsenate reductase and an ArsB arsenite efflux pump, *V. cholerae* lacks the latter and has an inactive ArsC. Therefore, *V. cholerae* does not reduce As^V^ or expel it out of the cell. From an evolutionary perspective, this could be a vestige of a functional ArsCB system that, after losing the ArsB component, has inactivated ArsC to prevent the generation of highly toxic As^III^. It remains to be investigated whether *V. cholerae* ArsC has undergone functional diversification to fulfill another role in the cell, perhaps related to its neighboring hypothetical genes (*vc2164-vc2167*).

Our screening identified a four-gene operon (*var operon*) conferring resistance to *V. cholerae* that encoded the ArsR repressor, the As^V^ permease ArsJ, a putative GAPDH (VarG) and a phosphatase (VarH). Interestingly, while most ArsR derepress transcription by sensing As^III^ (50), the lack of As^III^ production in *V. cholerae* suggests that ArsR probably senses As^V^ in this bacterium. Although more research is required to determine ArsR responses to As^V^ and As^III^ in *V. cholerae*, comparative sequence analysis between *V. cholerae* ArsR and that of As^III^ producing bacteria have revealed variations in conserved residues, including the E→K and C→A substitutions in the As^III^ binding motif (50) (Fig. S7A).

Although GAPDH enzymes are mostly known for their role in glycolysis, they have been functionally linked to diverse processes (51–53). A previous study reported that the GAPDH homolog to *V. cholerae’s* VarG detoxified As^V^ by generating 1As3PG from G3P. However, such conclusions were drawn from *in vitro* experiments that used commercial glycolytic GAPDH from rabbits instead of the specific As^V^-inducible GAPDH, and monitored As^V^ transport instead of direct 1As3PG detection (17). Our results, combining *in vitro* GAPDH activity assays and HRMS analyses from intracellular samples provided direct evidence that VarG is a GAPDH that preferentially uses As^V^ over P_i_ to synthesize 1As3PG. VarG role in *V. cholerae’s* As^V^ resistance adds to this growing list of GAPDH moonlighting functions.

Although the same study suggested that As^V^ is exported through ArsJ as 1As3PG (17), the more severe growth defect of *V. cholerae’s arsJ* compared to *varG* mutant suggests that 1As3PG does not play a major role in the export of As^V^ in this bacterium. Accordingly, the *arsJ* mutant accumulates intracellular As^V^ ca. 6 times more than the *varG* mutant. Even if we cannot wholly discount the transport of 1As3PG through ArsJ (as this compound can nonetheless be produced in the absence of VarG, likely by the constitutive glycolytic Gap activity), the fact that we have not detected extracellular 1As3PG or increased 3PG in cultures supplemented with As^V^ suggest that ArsJ expels free As^V^.

During glycolysis, the GAPDH enzyme catalyzes the simultaneous phosphorylation and oxidation of G3P to form 1,3 bi-phosphoglycerate (1,3-BPG), which is further converted into 3PG by phosphoglycerate kinase coupled to the transfer of the high-energy phosphate to ADP, making ATP. We reasoned that while the metabolic complexing of As^V^ into 1As3PG might decrease its toxicity by reducing As^V^ interaction with *V. cholerae*’s proteome (Fig4D), the formation of 1As3PG could be a dead-end product that would affect a major energy-producing pathway. In line with this reasoning, our results showed that VarH presents low phosphatase activity compared to canonical phosphatases. As low catalytic rate has been previously associated with enzymatic promiscuity (54), we hypothesized that the putative phosphatase VarH would function as an “arsenatase” that dissociates the 1As3PG complex to clear the glycolytic pathway and free As^V^ for its export through ArsJ. Indeed, inactivation of VarH results in increased intracellular levels of 1As3PG, which correlates with the accumulation of arsenate in the cell, thereby supporting the idea that ArsJ primarily exports free As^V^ in *V. cholerae*. Although synteny analyses suggest coevolution between ArsJ and VarG, the presence of VarH is rather exclusive to Vibrionaceae (Fig. S7B). It remains to be investigated whether the amino acid differences in ArsJ between VarG^+^ and VarG^-^ species explain the choice of transporting free or complexed As^V^.

It has been previously reported that arsenic induces the production of ROS in the cell (55–57). Our results show that in the presence of As^V^, *V. cholerae* induces oxidative stress responses: i.e., elevated levels of the hydrogen peroxide- tolerance determinant PrxA (58) and multiple proteins involved in the synthesis of cysteine. As cysteine-containing proteins are targets of ROS (59), and cysteine is also the building block of glutathione, stimulating its synthesis might help to alleviate oxidative damage provoked by As^V^. Additionally, we found that As^V^ directly or indirectly (via ROS) alters the stability of proteins implicated in energy- generating pathways such as TCA, fermentation, and respiration. Indeed, As^V^ accumulation in *var* mutant dissipated cellular membrane potential (a proxy of the cell’s energy status). The correlation between As^V^ accumulation with ROS and membrane depolarization exhibited by the *var* mutants suggests that oxidative stress and energy depletion might account for their growth impairment in As^V^.

*V. cholerae* is a pathogenic bacteria endemic to Bangladesh and India, regions where the concentrations of As^V^ in water and soil are the highest worldwide (60). Hence, *V. cholerae’s* superior resistance to As^V^ could have evolved as an adaptive strategy to thrive in As^V^-rich environments while endowing the cholerae pathogen with a fitness advantage over free-living and host-associated neighboring bacteria. Interestingly, despite the phylogenetic distance, heterologous expression of the *var* operon enhanced As^V^ resistance in all the pathogens assayed, demonstrating that this system has a high degree of versatility to work in combination with other arsenic detoxification systems.

Collectively, this work proposes a novel mechanism for As^V^ detoxification in *V. cholerae* that is entirely independent of As^V^-reduction and As^III^-extrusion. Our results support a model in which the formation of a transient As^V^-containing glycolytic intermediate cushions the impact of free As^V^ in the cell. While a metabolic sink might temporarily aid cells in enduring As^V^ toxicity, this strategy may exert a high cost in the long term as glycolysis is a major pathway in cell bioenergetics. We propose “dearsenylation” of 1As3PG by VarH provides a double benefit by decongesting any jam that may occur in glycolysis and freeing As^V^ for extrusion to the extracellular media by ArsJ.

The essentiality of the *V. cholerae’s var* operon to survive in environments with As^V^ (e.g., such as the host) suggests that this machinery may be considered a novel class of bacterial targets suitable for therapeutic intervention.

## ACKNOWLEDGMENTS.

This work was supported by the Knut and Alice Wallenberg Foundation (KAW), The Laboratory of Molecular Infection Medicine Sweden (MIMS), the Swedish Research Council and the Kempe Foundation. V.C.T and D.D.S. were supported by the National Institutes of Health/ National Institute of Allergy and Infectious Diseases (NIH/ NIAID) grant P01-AI83211 (Metabolomics Core) and R01- AI125588. The metabolomics analyses were performed at the University of Nebraska Medical Center Mass Spectrometry and Proteomics Core Facility administrated through the Office of the Vice-Chancellor for Research and supported by state funds from the Nebraska Research Initiative (NRI). We thank Erik Björk and Laurent Ouerdane for their help with the ICPMS analyses and for insightful discussions, J. J. Mekalanos for the *V. cholerae* C6706 Tn-mutant library and Inigo Ruiz and David Arranz for their technical support

## MATERIALS AND METHODS

### Bacterial strains and growth conditions

Strains used in this work are listed in Supplementary Table S2. *V. cholerae* strains used in this study are derivatives of the El Tor clinical isolate C6706. In addition to *V. cholerae*, in this study also were used the strains *Escherichia coli* K12 MG1655 and the *arsC*::Tn mutant strain from the Keio collection (21), *Enterohemorragic Escherichia coli* EHEC O157:H7, *Salmonella enterica serovar Typhimurium LT2*, *Citrobacter rodentium* DBS100, *Shigella flexneri* M90T, *and Yersinia pseudotuberculosis* YPIII. Strains were grown aerobically at 37°C in 15 ml tubes containing 3 ml of complete LB medium (10 g tryptone, 10 g NaCl, and 5 g yeast extract/L). To grow *Shigella flexneri,* TSB medium was used (Tryptone (Pancreatic Digest of Casein), 17 g Soytone (Peptic Digest of Soybean), 3 g Glucose, 2.5 g Sodium Chloride 5 g Dipotassium Phosphate 2.5). When used, casamino acids were supplemented at 1%. To study resistance to As^V^, 3 x 10^6^ overnight grown cells were inoculated into 200 µl of LB medium with or without As^V^ at indicated concentrations in 96-wells plates and OD values were measured every 10 minutes using a BioTek Epoch2 plate reader. Where appropriate, antibiotics were added to *V. cholerae* and *E. coli* cultures at the following concentrations: 200 (streptomycin, Sm), 100 (carbenicillin, Cb), 5 (Chloramphenicol, Cm) and 50µg/mL (kanamycin, Km).

Construction of plasmids to create *V. cholerae* mutants, overexpression constructs and transcriptional fusions.

*V. cholerae* mutants were created by allelic exchange with the suicide plasmid pCVD442 [48]. ca. 1kbp upstream and downstream DNA fragments flanking each coding regions were PCR-amplified with the primers listed in Table S3. Upstream and downstream DNA fragments were spliced together by overlapping PCR. The resulting ca. 2 Kb fragments were digested as indicated in table S3 and cloned into pCVD442. For allelic exchange of *V. cholerae arsC* by *E. coli arsC,* similar protocol was followed, but in this case the *V. cholerae* upstream and downstream DNA regions of *V. cholerae arsC* were spliced together with the *E. coli arsC* gene by overlapping PCR (see tables S2 and S3). Constructs were transformed into *E. coli* DH5α λpir for amplification. They were confirmed by sequencing, transformed into the *E. coli* donor strain SM10 λpir and conjugated for 6 hours at 37°C with *V. cholerae* C6706 by mixing equal volumes (1ml) of exponential phase cultures, and spot-plating. Single crossover *V. cholerae* were selected on LB plates with Sm and Cb. Re-streaked single colonies were then plated on salt-free LB agar containing 10% (w/v) sucrose and Sm. Colonies were streaked on carbenicillin plates to confirm loss of pCVD442 and then checked by PCR for successful deletion mutants.

Overexpression of *V. cholerae* and *E. coli* genes was carried out by using the arabinose inducible expression vectors pBAD33 and pBAD18, and the IPTG- inducible expression vector pHL100 (22) (see table S2 for specific constructs use for each enteropathogen). To construct each overexpressing plasmid, genes open reading frames including ribosome binding sites were PCR amplified (see primers used in table S3), double digested with indicated restriction enzymes and cloned into each plasmid. Constructs were transformed into the indicated strain by electroporation.

For construction of the transcriptional reporter fusions, gene promoter regions of target genes were amplified using the primers pair indicated in table S3. The PCR product was double digested with HindIII and EcoRI and then ligated into HindIII- EcoRI-digested pCB192N (23). Cloned plasmids were then transformed into *V. cholerae* by electroporation.

### Purification of recombinant VarG, VC2000, VC1041 and VarH

*V. cholerae* VarG, VC1041 and VarH proteins were overexpressed using the *E. coli* strain BL21 as 6×His-tagged enzymes from a pET28b vector. Overnight cultures were diluted into 250 mL LB broth and grown until OD_600_D=D0.5. Flasks were supplemented with 1 mM IPTG and shaking at 37° C for 2 h. Cells were pelleted, and resuspended in equilibration buffer (Tris 50 mM, pH 7.5, 50 mM NaCl, with protease inhibitor cocktail (Roche)) and disrupted by passaging once through a French press. Lysates were then centrifuged for 1 h (25,000 rpm, Beckman Coulter Avanti J26-XP centrifuge, JL-25.50 rotor) at 4°C. Nickel-NTA resin (0.5 ml resuspended in equilibration buffer) was then added to the supernatant, followed by incubation at 4°C in a rotating wheel. The lysate was separated from the resin by centrifugation for 1 minute at 3,220 g and the resin washed (5×10 ml) with washing buffer (equilibration buffer adjusted to 1 M NaCl) and eluted with 2ml of washing buffer containing 500mM imidazole. Fractions were subjected to SDS-PAGE and Coomassie Brilliant Blue staining for purity assessment. Protein concentration was quantified with a Bradford assay.

### Proteomic and thermal proteome profiling (TPP)

We performed thermal proteome profiling to determine protein abundance and thermal stability changes in WT, and *var* mutant strains upon exposure to As^V^ similarly to what was previously described (24, 25). Briefly, WT and mutant cells grown to an OD578 of 0.5 were incubated with 1 mM arsenate (or a similar volume of water) for 10 min. Cells were then washed with 10 ml PBS (for cells treated with arsenate, 1 mM arsenate were included in the wash step) and aliquoted to a PCR plate. Each aliquot was then exposed to a different temperature in the range 37°C-66.3°C for 3 min. Following a 3 min incubation at room temperature, cells were lysed with lysis buffer (final concentration: 50 μg ml^−1^ lysozyme, 0.8% NP-40, 1 × protease inhibitor (Roche), 250 U ml^−1^ benzonase, and 1 mM MgCl_2_ in PBS) for 20 min followed by three freeze–thaw cycles. Lysates were then prepared for analysis by mass spectrometry by digesting proteins using a modified sp3 protocol (26), labeling peptides with TMTpro (Thermo Fisher Scientific) and pooling samples from the same temperature together. These samples were then fractionated to six fractions with high pH fractionation and injected on an Orbitrap Q-Exactive Plus (Thermo Fisher Scientific) coupled to liquid chromatography (details on the run conditions and instrument parameters as in Mateus et al. (2020) (24, 25).

Mass spectrometry raw data was searched against the Vibrio cholerae FASTA file (UP000000584 downloaded from UniProt) using the Mascot 2.4 (Matrix Science) search engine and isobarquant (27). Protein abundance and thermal stability changes were determined using limma (28) using the same algorithm as in Mateus et al. (2020)(24).

### Phosphorylation state of VarG

To identify VC1069 GAPDH phosphorylation state*, V. cholerae* wild type and Δ*varH* strains carrying the overexpression vector pHL100 containing the *varG*-HIS tagged gene cloned were grown in the presence of As^V^ and with IPTG. Cells were pelleted, disrupted, load and run in SDS-PAGE gel. VarG-HIS bands were excised and diluted with NuPAGE LDS sample buDer (Invitrogen, Carlsbad, CA) mixed with10 mM dithiotreitol (DTT) and incubated at 56 °C for 20 min followed by addition of iodoacetamide (IAA) to a final concentration of 20 mM. The sample was loaded on 4-12% NuPAGE (Invitrogen). The electrophoresis was run in MOPS buDer at 180 V for 1 h. The gel was stained with Coomassie Brilliant Blue.

Bands of interest were cut from the SDS-PAGE. The bands were digested with trypsin, followed by identification using liquid chromatography–tandem mass spectrometry (LC–MS/MS). MS/MS spectra were searched with Proteome Discoverer 2.3 (Thermo Fisher Scientific) against the two sequences obtained for phosphate groups (UniProtKB). The precursor tolerance and fragment tolerance were set to 10 ppm and 0.05 Da, respectively. Trypsin was selected as enzyme, methionine oxidation, phosphorylation of serine, tyrosine, and tryptophan, and deamidation of asparagine and glutamine were treated as dynamic modification and carbamidomethylation of cysteine as a fixed modification (see supplementary methods for further information).

### Transposon-insertion sequencing (TIS) analysis

TIS was carried out essentially as previously described (29) but using selection plates containing LB medium with As^V^ 1 mM. In brief, ∼600,000 transposon mutants were generated by conjugation of *V. cholerae* C6706 with SM10λ*pir E. coli* carrying the Himar1 suicide transposon vector pSC189 (30). They were collected and their genomic DNA was pooled and analyzed on an Illumina MiSeq bench-top sequencer (Illumina, San Diego, CA). Insertion sites (which included 35% of TA sites) were identified as previously described (29), and significance was determined using Con-Artist simulation-based normalization as described (31). Results were visualized using Artemis (32).

### As^V^ resistance screen of arrayed transposon mutant library

A non-redundant transposon insertion library from the *Vibrio cholerae* strain C6706 (33) consisting of 3,156 insertion mutants was used to determine genetic determinants providing resistance to As^V^ in *V. cholerae*. *V. cholerae* WT and the library insertion mutants were first grown aerobically in 200 µl of LB in 96-well plates at 37°C for 24 hours. These cultures were then used as inoculum into 96- well plates containing 200 µl complete LB medium, in the presence of 1 mM of As^V^. The 96-well plates were incubated under agitation at 37°C for 24 hours and then resistance to As^V^ was evaluated by measuring optical density (OD_600_) using a BioTek Eon plate reader.

### Protein homology studies of As^V^ resistance determinants in enteric pathogens

DNA sequences of the *Vibrio cholerae arsR, varG, varH and arsJ var* genes and complete genomes sequences of the enteric pathogens: *Salmonella typhimurium* (PRJNA231221), *Citrobacter rodentium* (PRJNA261533), *Shigella flexeneri* (PRJNA553848), EHEC (PRJNA218110) and *Yersenia pseudotuberculosis* (PRJEB6403) were downloaded from NCBI database. Complete genomes of the pathogens were translated by Prodigal ver. 2.6.2 and homologous sequences to the *V. cholerae* Var proteins were queried into the enteric pathogens by BlastP ver. 2.9.0. Sequence homology data were represented in a heatmap diagram with Spearman correlation done by R ver. 3.6.1.

### β-Galactosidase activity determination

β-Galactosidase activity was measured through ONPG cleavage by the product of the *lacZ* reporter gene, and specific activity was calculated in Miller units (34).

In brief, cultures of *V. cholerae* carrying the pCB192N plasmid harboring promoter gene insertions (see table S2) were grown at 37°C overnight in 15 ml falcons tube containing 3 ml of LB complete medium. Overnight cultures were diluted 1:200 and supplemented with 1 mM of As^V^ and incubated at 37°C until they reached mid-log phase (OD_600_ 0.4-0.7). A 100 µl aliquot of three different subcultures were collected, cells were permeabilized and assayed in triplicate for each strain as previously described (34).

### Determination of intracellular and extracellular As^V^ and As^III^ in *V. cholerae*

To determinate concentrations of As^V^ and As^III^ in supernatants of *V. cholerae* and *E. coli,* bacterial cells were grown in the presence of As^V^ for 5 hours and then pelleted. Supernatants of each strain were speciated by high-performance liquid chromatography (HPCL) (Agilent Infinity II 1290) using a C18 column reverse- phase (130 Å, 1.7 μm, 2.1 mm by 150 mm; Waters, USA). Elution conditions used were flow rate 1 ml min^-1^; temperature 25 °C; isocratic elution in 5 mM tetrabutylammonium hydroxide, 5 % methanol (v/v) and 3 mM malonic acid. Identification and quantification of As^V^ and As^III^ was performed by inductively coupled plasma mass spectroscopy (ICP-MS) (Agilent 8900 ICP-MS Triple Quad) by comparison to standards of known concentration and peak integration.

To determine intracellular concentrations of As^V^ and As^III^ in *V. cholerae,* the pelleted cells were washed in complete LB medium to remove traces of arsenic, then resuspended in 500 µl of H_2_O. Glass beads were adding to the resuspension and cells were lysed by bead-beating. The resulting extract was centrifuged, and As^V^ and As^III^ from those supernatants were analyzed by ICP-MS as previously described. Samples were analyzed in triplicate.

### LC-MS/MS analysis using QTRAP6500+

To measure 3-phosphoglycerate (3PG), *V. cholerae* cells grown in the presence of 5 mM of As^V^ were pelleted and used for LC-MS/MS analysis. Bacterial pellets were resuspended in 1 ml ice-cold 60% ethanol containing 2 μM ribitol as the internal control. The cells were mechanically disrupted using a bead homogenizer set to oscillate for 3 cycles of 30 sec each at 6800 rpm with a 10 sec pause between each cycle. Cell debris was separated at 13,000 rpm. An aliquot (200µl) of the supernatant containing intracellular metabolites was vacuum dried and resuspended in an equal volume of 7.5 mM ammonium bicarbonate solution before LC-MS/MS analysis.

A triple-quadrupole-ion trap hybrid mass spectrometer, QTRAP® 6500+ (Sciex, USA) connected with a Waters ultra-performance liquid chromatography I- class (UPLC) system was used for metabolite analysis. The chromatographic separation was performed on an XSELECT HSS XP column (150Dmm ×D2.1Dmm ID; 2.5 µm particle size, Waters, USA) using a binary solvent system at a flow rate of 0.1 ml/min. Mobile phase A was composed of 7.5 mM ammonium bicarbonate in LC-MS grade water and mobile phase B was 100% LC-MS grade methanol. The column was maintained at 40°C, and the autosampler temperature was set to 7°C. The A/B solvent ratio was maintained at 100/0 for 2Dminutes, followed by a gradual increase of solvent B to 95% for 2 minutes. The solvent B was maintained at 95% over the next 5 minutes. The gradient was reduced to 100% solvent A within 0.5 min and column was equilibrated for 5.5Dminutes before the next run. The needle was washed with 1.2 ml of a strong wash solution containing 100% LC-MS-grade acetonitrile followed by 1 ml of a weak wash solution comprised of 10% aqueous methanol before each injection. The injection volume was 5µl. The QTRAP® 6500+ (SCIEX) was operated in negative as well as positive ion mode for targeted quantitation in Multiple Reaction Monitoring (MRM). MRM parameters for each analyte are listed in Table 1. The electrospray ionization (ESI) parameters used are as follows: electrospray ion voltage of -4500V in negative and 5500V in positive mode, source temperature of 400°C, curtain gas of 35, and gas 1 and 2 of 40 psi each, respectively. The compound-specific parameters such as declustering potential (DP) and collision energy (CE) were optimized for each compound using manual tuning. These values are listed in Table 1.

**Table 1:** MRM transitions, declustering potentials and collision energies of metabolites

| Analyte | Q1 (m/z) | Q3 (m/z) | DP (V) | CE (V) |
| --- | --- | --- | --- | --- |
| 3-phospho glycerate | 187.2 | 98.9 | +60 | 19.2 |
| Ribitol | 150.8 | 88.9 | -60 | -14.2 |

### Detection of 1As3PG

1As3PG was identified using a Thermo Orbitrap Exploris 480 high resolution mass spectrometer. The chromatographic method used for this analysis was the same as that described for the LC-MS/MS performed using the QTRAP6500+. The MS data were acquired in both positive and negative polarity within mass range of 140- 400 Da. The precursor ion resolution was maintained at 240,000, whereas the product ion resolution at 120,000. Deprotonated [M-H]^-^ ion of 1As3PG was detected at m/z 304.8696 Da compared to the actual mass of 304.8674 Da, thus Δm = 7.149 ppm. The fragmentation pattern of 1As3PG could not be further resolved under these conditions.

### Proteins structural and topological analysis

Protein alignments were performed by using Vector NTI. Prediction of VarH secondary structure was performed by using Phyre2 (http://www.sbg.bio.ic.ac.uk/phyre2/html/page.cgi?id=index). Structural prediction of the VarG and VarH proteins was performed with AlphaFold2 on the CoLabFold publicly accessible interface (35) (https://colab.research.google.com/github/sokrypton/ColabFold/blob/main/AlphaFold2.ipynb). Sequences were modeled as monomers using mmseqs2 for multiple sequence alignment. Studies on proteins topological similarity was carried out by using TM-score (https://zhanggroup.org/TM-score/).

### Competition assays

Aerobic overnight cultures of V. cholerae C6706 wild type lacZ^-^ and Enterohemorragic Escherichia coli EHEC O157:H7 lacZ^+^, Salmonella enterica serovar Typhimurium LT2 lacZ^+^, Citrobacter rodentium DBS100 lacZ^+^, Shigella flexneri M90T lacZ^+^, and Yersinia pseudotuberculosis YPIII lacZ^+^, were collected and 1 x 10^7^ cells co-inoculated at 1:1 ratio in 15 ml tubes containing 3 ml of complete LB medium without or with 10 mM As^V^. After 8 hours of incubation at 37°C, an aliquot was diluted and plated on LB agar plates containing X-Gal to enumerate wild type and competing strains. Competitive indices (CIs) were determined by dividing the ratio mutant or non-wild type strain to C6706 wild type colonies by the ratio in the inoculum. To calculate CIs among V. cholerae strain C6706 against a different V. cholerae strain against enteric pathogens (Fig. 1B), CIs obtained after the As^V^ challenge were normalized respect CIs obtained after competition in the absence of As^V^.

### *in vitro* GAPDH and phosphatase enzymatic assays

The GAPDH activity assay was performed in accordance with manufacturer’s instructions (MAK27; Sigma-Aldrich) but using P_i_ or As^V^ as the GAPDH substrate. 10 µg of purified *V. cholerae* VarG and VC2000 enzymes were used in the assay. This assay quantifies GAPDH activity by measuring enzymes capacity to reduce NAD^+^ to NADH during the conversion of glyceraldehyde 3-phosphate to 1,3- biphosphoglycerate. NADH produced in this reaction results in a colorimetric (OD_450_) product proportional to the GAPDH activity of the enzyme. GAPDH activity is reported as nmole/min/mL = milliunit/mL. One unit of GAPDH is the amount of enzyme that will generate 1.0 mmole of NADH per minute at pH 7.2 at 37 °C.

Phosphatase activity was quantified using Cayman’s Phosphatase Colorimetric assay kit and 5 µg of purified *V. cholerae* VarH, C113G R119G VarH, VC1041 (positive control) and BsrV (negative control) enzymes. The assay uses *p*-nitrophenylphosphate (pNPP) as substrate for phosphatase enzymes. Phosphatase enzymes dephosphorylate pNPP, which deprotonates its phenolic OH-group by increasing the pH of the reaction, deprotonated pNP yields an intense yellow color detected at OD 405 nm. Enzymatic activities were calculated as µmol of phosphate released per min/ml.

### Flow cytometry analysis

*V. cholerae* WT and indicated *var* mutant strains were grown for 5 hours in the presence of 1 mM As^V^, then Aliquots (1 ml) from each culture were pelleted, washed once, and resuspended in PBS (100 µl) to measure membrane potential and ROS.

To measure *membrane potential* in *V. cholerae* cells, we used the BacLight bacterial membrane potential kit (Invitrogen). *V. cholerae* samples resuspended in 100 µl PBS were incubated for 15 min at 20 °C in the dark with the dye 30 μM 3,3’-diethyloxacarbocyanine iodide (DiOC_2_). Next, cells were washed twice with PBS and resuspended in 2 ml of PBS, then fluorescence emitted from treated cells was measured with a Bio-Rad S3e cell sorter at an excitation wavelength of 488Dnm and an emission wavelength of 525/30 nm (for green) or 655 nm (for red).

*ROS measurements* were performed using the CellROX™ Green Reagent, for oxidative stress detection (ThermoFisher). Washed *V. cholerae* cells were supplemented with CellROX® Reagent at a final concentration of 5 μM and incubate them for 30 minutes at 37°C. Next, cells were washed twice with PBS and resuspended in 2 ml of PBS, then fluorescence emitted was measured with a Bio-Rad S3e cell sorter at an excitation wavelength of 488 nm and an emission wavelength of 525 nm.

### Statistics and reproducibility

Statistical significance was assessed with the Student’s *t*-test where indicated. A p-value of less than 0.05 was considered statistically significant. Assays were performed with three biological replicates unless otherwise indicated.

## SUPPLEMENTARY INFORMATION

**Fig. S1 Growth curves (OD600) of V. cholerae WT in the presence of AsV.** *V. cholerae* WT was grown in LB medium in the presence of 10, 20 and 30 mM As^V^. Data are the mean of three biological replicatesD±Ds.e.m.

**Fig. S2 Characterization of V. cholerae ArsC. A** Schematic of the *arsC*- containing operons from *E. coli* K12 and *V. cholerae,* and protein sequence alignments. Black arrows depict ArsC catalytic cysteine and arginine residues implicated in As^V^ reduction to As^III^. **B** Growth curves (OD_600_) of *V. cholerae* WT, and two derivative strains where *arsC* has been replaced by *E. coli’s arsC* (red, *arsC*^ec^), and *arsC*^ec^ strain where *arsB* is overexpressed in trans (blue, *arsC^ec^ + arsB^ec^*) (see bottom panel, and material and methods for further details). Cultures were grown in LB medium in the presence 10 mM As^V^. Bottom panel: Explanatory diagrams illustrating the expected output from each construct. Data are the mean of three biological replicatesD±Ds.e.m.

**Fig. S3 Transcriptional analysis of the var operon.** β-galactosidase activity from *V. cholerae* WT and *arsR* mutant (Δ*arsR*) strains carrying DNA regions upstream of *arsR, varG, varH* and *arsJ* genes fused to the *lacZ* reporter gene. Each DNA region fused to *lacZ* is depicted by arrows. Cultures were grown in LB medium in the presence or absence of 10 mM As^V^. Data are the mean of three biological replicatesD±Ds.e.m.

**Fig. S4 V. cholerae VarG presents a high degree of similarity to the glycolytic GAPDH Gap. A** Sequence alignments of the *V. cholerae* VarG and the glycolytic GAPDH proteins using Vector NTI software. **B** Left: Predicted structures of VarG using AlphaFold2 (https://colab.research.google.com/github/sokrypton/ColabFold/blob/main/AlphaFold2.ipynb). The region of the protein structure depicted in blue colour indicate the least error in prediction. Right: VarG and Gap structures overlapping using TMscore (https://zhanggroup.org/TM-score/). TM-score value of 1 indicates 100% structural identity among proteins.

**Fig. S5 Quantification of intra- and extracellular metabolites from *V. cholerae*.** Representative plots from identification of 1As3PG and 3PG from intracellular and extracellular samples from *V. cholerae* WT and Δ*varG* cells were grown with and without 1 mM As^V^. The compounds were separated by UPLC using an XSELECT HSS XP column. 1As3PG was identified using a Thermo Orbitrap Exploris 480 high-resolution mass spectrometer and 3PG using a triple-quadrupole-ion trap hybrid mass spectrometer QTRAP® 6500+ (Sciex, USA) connected with a Waters ultra-performance liquid chromatography I-class (UPLC) system.

**Fig. S6 VarH characterization. A** Sequences alignment of the secondary structure of the VarH protein with the crystal structure of the human Kap PTP phosphatase using the software Phyre2 (www.sbg.bio.ic.ac.uk/phyre2). Human Kap structure matched VarH with 100% confidence and 99% coverage. Amino acid sequences within the black box (from the amino acid 113 (H) onwards) indicate the conserved catalytic loop (C(x)_5_R) of protein tyrosine phosphatases PTP. Green and black boxes indicate deletions and insertions, respectively, respect the template sequence. **B** Predicted structures of VarH with AlphaFold2 (https://colab.research.google.com/github/sokrypton/ColabFold/blob/main/AlphaFold2.ipynb). The regions of the protein structure depicted in blue colour indicate no error in prediction. Right: overlapping between VarH protein structure obtained with AlphaFold2 and crystal structure of the Kap PTP protein using TMscore (https://zhanggroup.org/TM-score/). TM-score value of 1 represents 100% structural identity among proteins. **C** SDS-PAGE gel depicting the VarG His-tag purified protein from *the V. cholerae* WT and *varH* mutant strains (see material and methods for details). VarG proteins were excised from the gel and its phosphorylation state was quantified by LC–MS/MS. **D** Phosphorylation state quantification of VarG from *V. cholerae* WT and *varH* mutant strains. S204, serine amino acid from VarG in position 204. Y108, tyrosine amino acid from VarG in position 108. **E** Representative plots from identification of 1As3PG and 1AsG from intracellular samples of *V. cholerae* WT and Δ*varH* cells grown with and without 1 mM As^V^. The compounds were separated by UPLC using a XSELECT HSS XP column. The detection of 1As3PG and 1AsG were perfomed using a Thermo Orbitrap Exploris 480 High Resolution Mass Spectrometer. Data are the mean of three biological replicatesD±Ds.e.m.

**Fig S7 AsIII binding site of ArsR is not conserved in *V. cholerae* and *varH* is specific of the Vibrionaceae family. A** ArsR amino acids sequences alignment from *V. cholerae* and representative enteropathogenic bacteria using Vector NTI software. ArsR cysteine-containing As^III^-binding motives are depicted in the lower panel with cysteine residues labeled in red color. **B** Genomic organization comparison of the *var* operon in the Vibrionaceae family and representative bacteria by using the Patric software (https://www.patricbrc.org/). The presence of *varH* is depicted in green color and under dotted rectangle.

Table S1. Complete list of genes identified under-represented by TIS in the presence of 1 mM As^V^ with thresholds of fold change <0.1 and an inverse p-value >75

Table S2. List of strains and plasmids used in this study.

Table S3. List of primers used in this study.

